# Epstein-Barr virus nuclear antigen proteins deploy diverse mechanisms to bind the human genome

**DOI:** 10.1101/2022.07.18.500442

**Authors:** Md. Redwanul Haque, Abdur Rashid Tushar, Minjun Park, Chang In Moon, M. Sohel Rahman, Md. Abul Hassan Samee

**Author notes:** These authors contributed equally to the work (Md. Redwanul Haque, Abdur Rashid Tushar).

## Abstract

Viral transcription factors (vTFs) are known to bind the human genome and impact critical biological processes. However, a vTF can infect multiple cell-types, and the extent to which they deploy the same mechanisms for DNA binding across different cell-types is poorly understood. We used the Epstein-Barr virus Nuclear Antigen Proteins 2 and 3 (EBNA2 and EBNA3) to address this gap, the two widely studied vTFs associated with cancers, as case studies. We analyze multiple ChIP-seq datasets of these vTFs from different human cell-lines using a state-of-the-art convolutional neural network model. We show that each vTF uses both cell-type specific and cell-type independent cofactors to bind the human genome. Interestingly, each vTF requires ∼10 cofactors consistently across all binding peaks. Such dependency on a large number of cofactors is in contrast to human TFs since human TFs typically co-bind with 2-3 cofactors. Our *de novo* motif finding approach also reveals novel sequence motifs at the ChIP peaks of these vTFs. These critical signals were missed by previous studies doing enrichment analysis using known human TFs. Finally, we find that although a vTF impacts similar biological processes across cell-types, it also impacts distinct biological processes in a cell-type specific manner. Our study provides a roadmap for integrative analysis of vTF binding and pinpointing their diverse mechanisms of targeting the human genome across different human cell-types.

## Introduction

Humans are infected by over 200 viral species [1]. A wide range of cytopathic effects is induced when human cells are infected by viruses. Some viruses may control host gene expression or genome replication or transmission to other cells and lead to cytopathic effects. The genome of viruses encodes multiple proteins that cause these effects. Viral transcription factors (vTFs), cofactors, and other regulators of gene expression are some of these proteins. These proteins can control both the expression of viral and host genes and so are core to the pathogenesis of virus-induced diseases in humans. These proteins are referred to as vTF which can modulate gene transcription through direct or indirect interactions with nucleic acids.

Epstein-Barr virus (EBV) is one of the viral species which is associated with many diseases. It was discovered in Burkitt’s lymphoma (BL) biopsy cell culture in 1964 [2] and was later associated with it [3]. More studies have revealed the causative role of EBV in the development of various diseases and cancers including Hodgkin’s lymphoma [4], mononucleosis [5], multiple sclerosis [6–8], inflammatory bowel disease (IBD) [9], rheumatoid arthritis (RA) [10], and SLE [11]. EBV contains <u>E</u>pstein-<u>B</u>arr <u>n</u>uclear <u>a</u>ntigen proteins (EBNAs) that have several groups within them. In our study, we focus on the two well-studied families, EBNA2 and EBNA3.

A strong line of research has revealed how the EBNA2 and EBNA3 proteins molecularly associate with human diseases. EBNA2 is a transactivator protein that is expressed in latently infected B lymphocytes. It takes part in the regulation of latent vTFs and plays a role in the immortalization of EBV infected cells [12][13]. EBNA2 controls the processes of immortalization of EBV-infected B cells by altering the expression levels of human genes [14]. It has been suggested that EBNA2 mediates some of this regulation through interactions with human TFs such as EBF1, RBPJ, and SPI1 (also known as PU.1) [15]. The EBF1 protein, for example, binds to and opens chromatins that are occupied by EBNA2-RBPJ complexes [16]. Likewise, EBNA3C has been reported to impact gene regulation by interacting with RBPJ [17]. In a study, EBNA2/3B/3C are reported to target the genes associated with B cell activation and B cell transcription related biological processes that cause rewiring of B cell function and large scale modulation of B cell transcriptional programs [15]. In another study, it is said that these vTFs target genes associated with inflammation, apoptosis, and signaling pathways (P38, MAPK, p53, Toll-like receptor, and JUN Kinase) which suggests that transcriptional programs that rewire cellular responses to various stimuli and promote cell survival can be initiated by EBV vTFs [18]. In the same study, it is claimed that EBV might target the important developmental Wnt pathway [18].

Despite these studies on the EBNA proteins’ roles in pathogenesis and the biological processes impacted, there are critical knowledge gaps regarding the mechanisms of how these proteins target specific genomic loci in the human genome. To what extent do the sequence signatures, i.e., motifs, at their binding loci remain consistent across cell-lines? Do they target unique sequences where human TFs do not bind? Do they require cobinding factors (cofactors) and do the cofactors remain the same across cell-lines? Do they impact the same biological processes across cell-lines? These are some of the questions that we address in the present study.

We used the ENCODE consortium’s recommended steps to reprocess three ChIP-seq datasets of EBNA2 and four datasets of EBNA3. Using state-of-the-art deep learning methods to model these ChIP-seq data, we found that both proteins have cell-line specific and cell-line independent binding motifs and cofactors. Our models also revealed some unique motifs that do not match the motifs of known human TFs even at very flexible thresholds for motif similarity. Notably, the sequence specificity of these “novel” motifs is comparable to other human TF motifs. Furthermore, our analyses revealed that each vTF requires ∼10 cofactors consistently across all binding peaks in the human genome. This intriguing dependency of vTFs on a large number of cofactors is in contrast to human TFs since human TFs typically co-bind with 2-3 cofactors. Finally, we find that both common and cell-line specific biological processes and pathways are impacted by EBNA2 and EBNA3. Overall, our study offers the first comprehensive investigation into the cell-line specificity of DNA binding mechanisms deployed by EBNA2 and EBNA3.

## Results

### A Convolutional Neural Network (CNN) model of DNA binding specificity of EBNA proteins

CNN models have shown superior performance over traditional machine learning models for classification modeling of genomic sequence data [19]. However, revealing the class-specific sequence motifs from these models is not straight-forward. In order to build CNN models that are both accurate and easy-to-interpret, we used the recently proposed MuSeAM model (Fig 1A). MuSeAM is a custom CNN model shown to be more interpretable than conventional CNN for sequence analysis [20]. We used the model for the prediction of DNA binding specificity of EBNA2 and EBNA3 vTFs. We also benchmarked MuSeAM against conventional CNN (Fig 1B) models.

**Fig 1.**
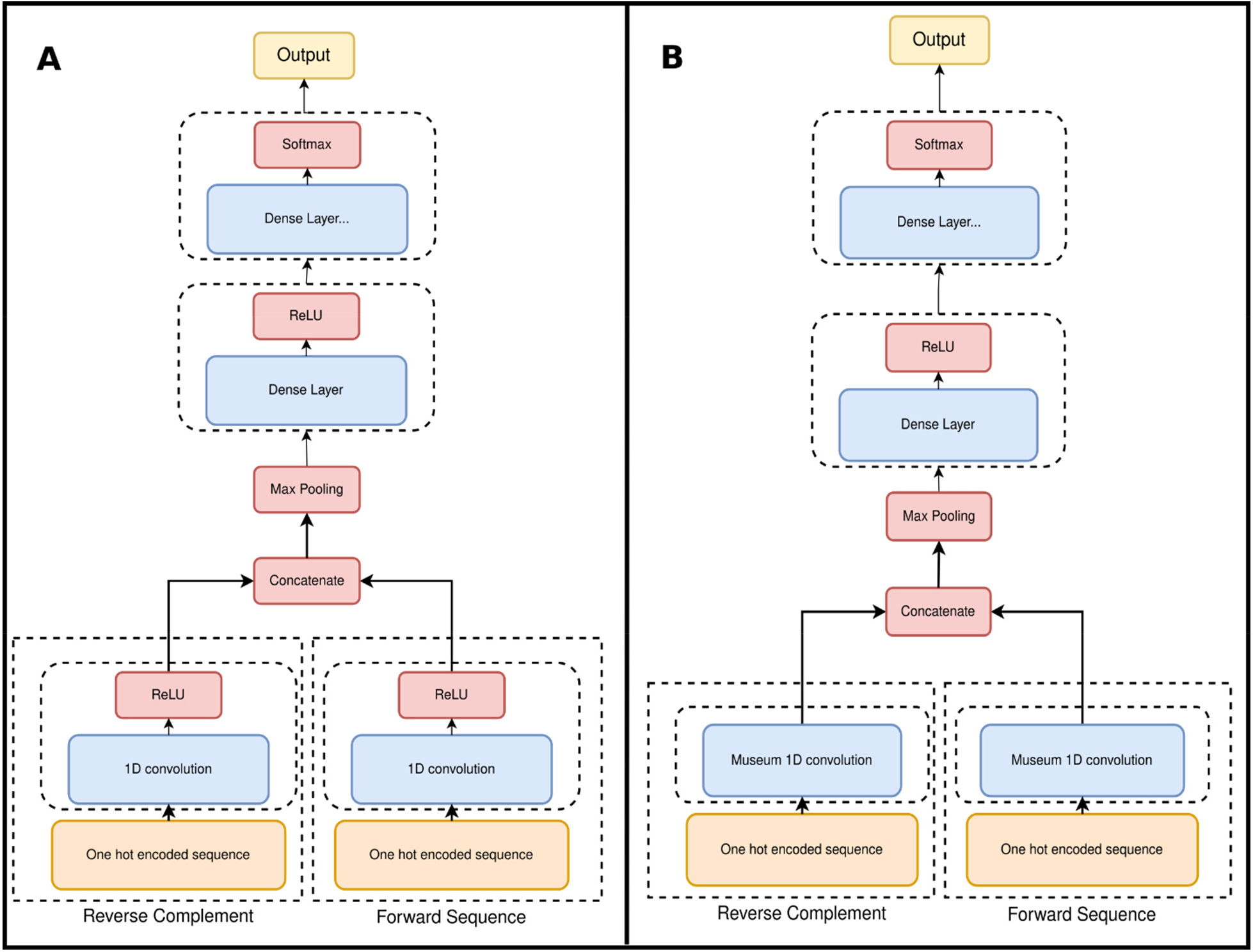
Model Architecture. (A) The MuSeAM model. One hot encoded forward and reverse complement sequences are input into the model which are followed by a MuSeAM 1D convolution layer. The motifs are obtained from this convolution layer. Then the outputs of the convolutions are concatenated and followed by a max-pooling layer. Then a fully connected layer is applied followed by a ReLU activation. Finally, the classification layer is applied which is a fully connected layer of two features followed by a softmax. (B) The conventional CNN model almost has the same architecture as (A) except that the MuSeAM convolution layer is replaced by conventional 1D convolution followed by a ReLU activation.

### MuSeAM predicts EBNA2 and EBNA3 binding sites with high accuracy

We have modeled seven Epstein-Barr-Virus (EBV) ChIP-seq datasets in this study. These datasets were mentioned in a recent review on vTFs by Liu et al. [18]. Of these, three datasets feature EBNA2 binding in the IB4, GM12878, and Mutu3 cell-lines (peak counts of 1344, 1322, and 5288, respectively). The other four datasets are EBNA3A, EBNA3B, and EBNA3C binding in the LCL cell-line and EBNA3ABC in the Mutu3 cell-line (peak counts of 21, 1176, 72, and 1764 respectively). Given that the peak counts for EBNA3A and 3C were very low in LCL, we created a combined dataset by merging the EBNA3A, EBNA3B, and EBNA3C peaks in the LCL cell-line and named it EBNA3ABC-LCL. All datasets were generated from raw ChIP reads using the uniform ChIP data processing pipeline recommended by the ENCODE consortium [21]. As negative sequences for the classification model, we generated null sequences using the gkmSVM R package [22]. Fitting both MuSeAM and the conventional CNN models under 10-fold cross validation showed the models are nearly identical (Table 1). However, as MuSeAM offers an extra edge in terms of the first layer kernels being directly interpretable as sequence motifs, we used MuSeAM and its motifs throughout our analyses. Through hyperparameter searching, we fixed 16 kernels of length 14 on all the three EBNA2 datasets and the EBNA3ABC-Mutu3 dataset, and 48 kernels of length 8 on the EBNA3ABC-LCL dataset.

**Table 1.** Benchmarking metrics of MuSeAM and conventional CNN.

| Viral Protein | Cell-line | Accuracy |  | AUROC |  |
| --- | --- | --- | --- | --- | --- |
|  |  | MuSeAM | Conventional | MuSeAM | Conventional |
| EBNA2 | Mutu3 | 0.867 | 0.867 | 0.943 | 0.940 |
|  | IB4 | 0.916 | 0.910 | 0.972 | 0.974 |
|  | GM12878 | 0.854 | 0.872 | 0.930 | 0.947 |
| EBNA3ABC | LCL | 0.874 | 0.819 | 0.939 | 0.903 |
|  | Mutu3 | 0.852 | 0.863 | 0.93 | 0.938 |

For each viral protein the trained MuSeAM model of one cell-line was used to predict on datasets of other cell-lines. Cross results for EBNA2 are mentioned in the S2 Table. For EBNA2, the model trained on one cell-line performs fairly well on the other ones too. Each model is best on its own dataset, so there must be some unique target genomic loci in each cell-line. But as the models are much better than random on other cell-lines, we can say that there are also common target genomic loci across cell-lines.

Cross results for EBNA3 are mentioned in the S2 Table. Same as EBNA2, we see for EBNA3 that models perform best on their own cell-lines, but can also predict well on the other cell-lines, showing signs of both similarities and uniqueness of target genomic loci across cell-lines.

### Motifs discovered from EBNA ChIP-seq data suggest extensive co-binding of EBNA and human proteins

Using MuSeAM kernels, we predicted EBNA2 motifs and used the Tomtom [23] tool to match the motifs with the known human libraries of human TFs. Five out of 16 predicted motifs for EBNA2 in the IB4 cell-line show strong similarity with known human motifs, namely the TFs COE1 (EBF1), SUH (RBPJ), ZNF18, ZN770, and ZN121 (Fig 2).

**Fig 2.**
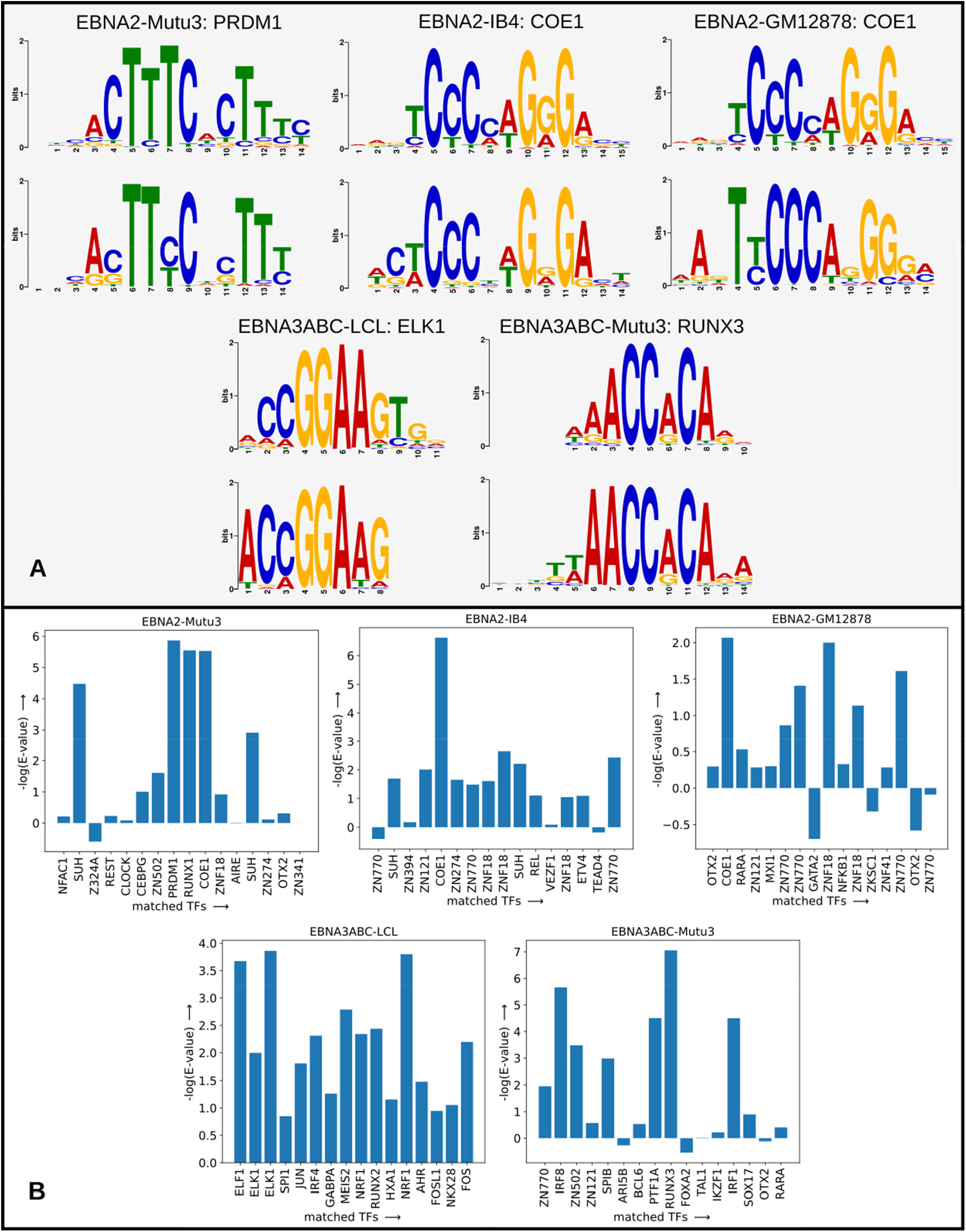
Results obtained from Tomtom and corresponding statistics. (A) The motif with the highest E-value with its best match is shown for each cell-line. For each pair, the motif on top is a known human TF and on the bottom is the motif predicted by our model. (B) Along the X-axis are names of the best matches for each motif in each cell-line with known human TFs found using Tomtom (only the top 16 motifs sorted using the E-values of their best match were taken for the EBNA3ABC-LCL cell-line). The corresponding E-values of the best matches are shown along the Y-axis in the negative logarithmic scale.

For the EBNA2 motifs in the GM12878 cell-line, we found strong similarities with four out of 16 motifs, namely the TFs COE1 (EBF1), SUH, ZNF18, and ZN770. Finally, five out of 16 EBNA2 motifs in the Mutu3 cell-line showed strong similarity with known human motifs, namely the TFs PRDM1, RUNX1, COE1 (EBF1), and SUH (RBPJ). Across all cell-lines, the EBNA2 motifs discovered by MuSeAM matched the EBF1 and SUH (RBPJ) motifs with significant Tomtom E-values (Fig 2B). This conforms to prior reports on EBNA2 binding sites being highly enriched for these motifs [24] [25]. Also, another study on TF motif enrichment analysis revealed that motifs for the TFs EBF1, BCL11A, RBPJ, Ets are enriched, in this order, in the binding sites of EBNA2 [18], our model found motifs that match these TFs as well. But in that analysis PRDM1 motif is not mentioned as a significant binding site. But our model predicts a motif with strong match with PRDM1 in the Mutu3 cell-line (Fig 2). This is supported by a study where it mentioned the association of PRDM1 with the EBV lymphomagenesis [26].

Our analysis of the MuSeAM motifs for EBNA3 showed similar matches with motifs for known human TFs. Of the 48 predicted motifs for EBNA3ABC-LCL cell-line seven show strong similarity with known TFs, including the TFs ELK1, NRF1, ELF1, MEIS2, and RUNX2. Of the 16 predicted motifs for the EBNA3ABC-Mutu3 cell-line 6 show strong similarities with known TFs, including the TFs RUNX3, IRF8, IRF1, PTF1A, and ZN502 (Fig 2). Our model also found motifs that match motifs in the previously mentioned study on TF motif enrichment analysis which revealed that motifs for the TFs ISRE, Ets, BCL11A, RUNX are enriched in the binding sites of EBNA3 [18].

### Novel motifs discovered in the EBNA2 and EBNA3 ChIP-peaks

Based on both Tomtom E-value and visual inspection, some motifs learned by MuSeAM are very poor matches to known human TF motifs. We call these motifs *novel* (explained in Methods section) for the respective viral TFs. We found novel EBNA2 motifs in the GM12878 and Mutu3 cell-lines, and for EBNA3ABC in LCL (Fig 3A).

**Fig 3.**
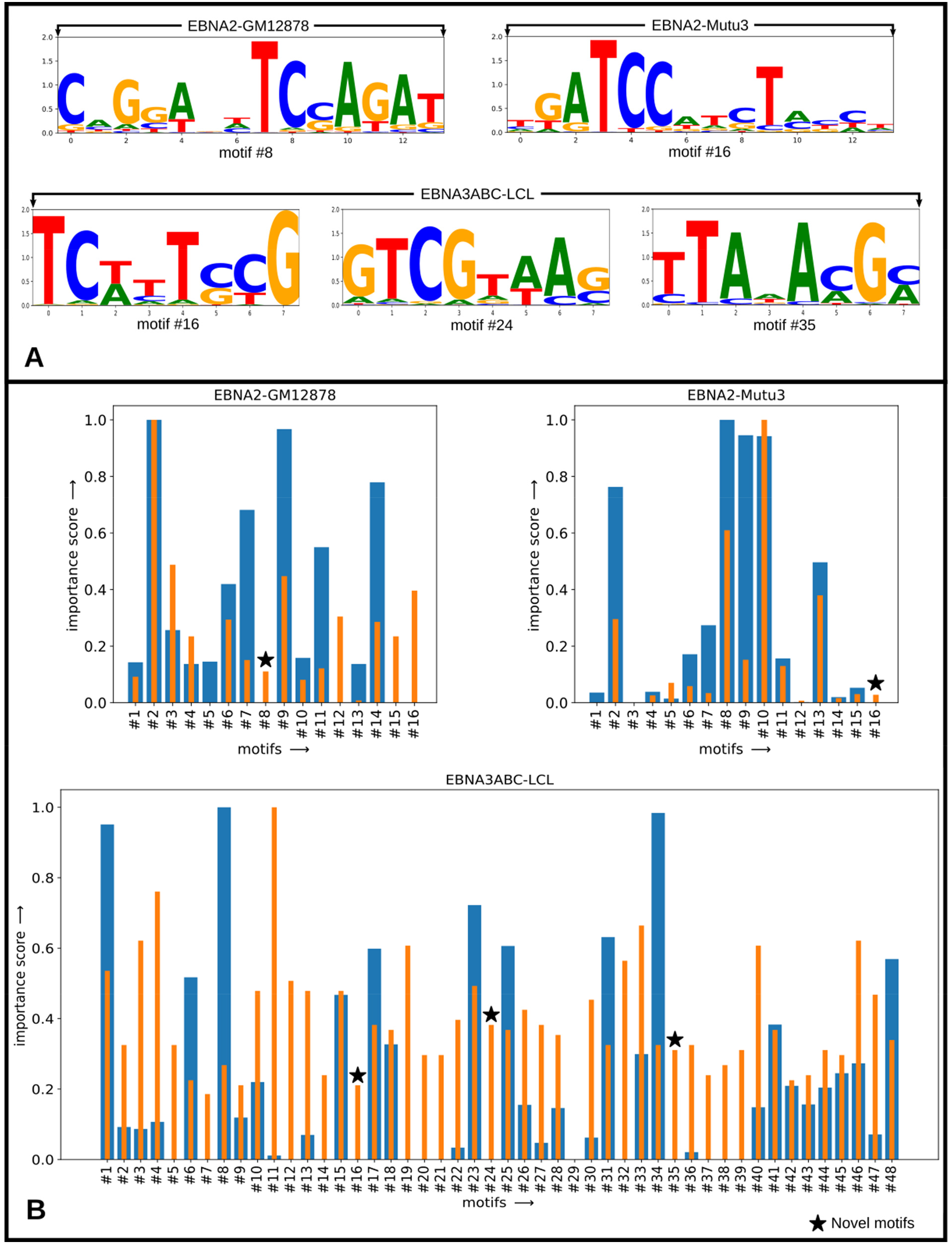
Novel motifs and their importance scores. (A) One novel motif found for the EBNA2-GM12878 cell-line, one for the EBNA2-Mutu3 cell-line, and three for the EBNA3ABC-LCL cell-line. The height of each nucleotide character is proportional to their respective information content. These motifs were drawn using standard Tomtom convention, i.e: without considering background probabilities. (B) Each motif of the three cell-lines where novel motifs were found is given two importance scores, both in range [0, 1]. The blue bar for each motif represents the importance score derived by normalizing the negative logarithm of the E-values of the best matches found for all motifs using Tomtom. The orange bar for each motif represents the importance score derived by normalizing the accuracy drop suffered by a model when that motif is removed.

Even at an E-value cutoff of 1.0, Tomtom could not detect a matching human TF for these motifs. Importantly, these motifs are not degenerate (non-specific): the average Shannon’s information content per nucleotide position in these motifs is 1.396 bits for the EBNA2-GM12878 motif, 1.381 bits for the EBNA2-Mutu3 motif, and for motif#16, motif#24 and motif#35 of EBNA3ABC-LCL it is 1.598 bits, 1.639 bits and 1.429 bits respectively. Compared to 1.27 bits of information content per position in human motifs in the TRANSFAC library [27], these motifs are specific and Tomtom’s failure to find their match with human TF motifs is likely not due to degeneracy. In order to see if the novel motifs are important in the prediction capability of the model, we ranked each motif by individually removing it from the model and using the adjusted model to predict on the dataset again. Each motif was then given an importance value between 0 and 1 by normalizing the accuracy drop of all motifs (Fig 3B).

### Predicted motifs are common to all peaks of a dataset

We next checked if a vTF’s peaks can be partitioned into groups based on the occurrence of different motifs. Thus, we calculated the matching score of each motif on each peak of a dataset (Methods). We did not find any motif favoring certain groups of peaks. Instead, consistently across all peaks, the more important a motif the more is its matching score (Fig 4). This characteristic of vTFs using >10 cofactors consistently across their ChIP-seq peaks is in striking contrast with how human TFs utilize cofactors. First, prior analysis has shown that human TFs use fewer, typically 2-3 cofactors, at their ENCODE ChIP-seq peaks [28]. Secondly, at a genomewide scale, human TFs do not generally require all cofactors at their DNA binding loci [29]. Overall, powered by *de novo* motif finding, our analyses revealed an intriguing dependency of vTFs on human cofactors for binding the human genome. This signal was missed by prior studies performing conventional motif enrichment analysis at vTF ChIP-peaks.

**Fig 4.**
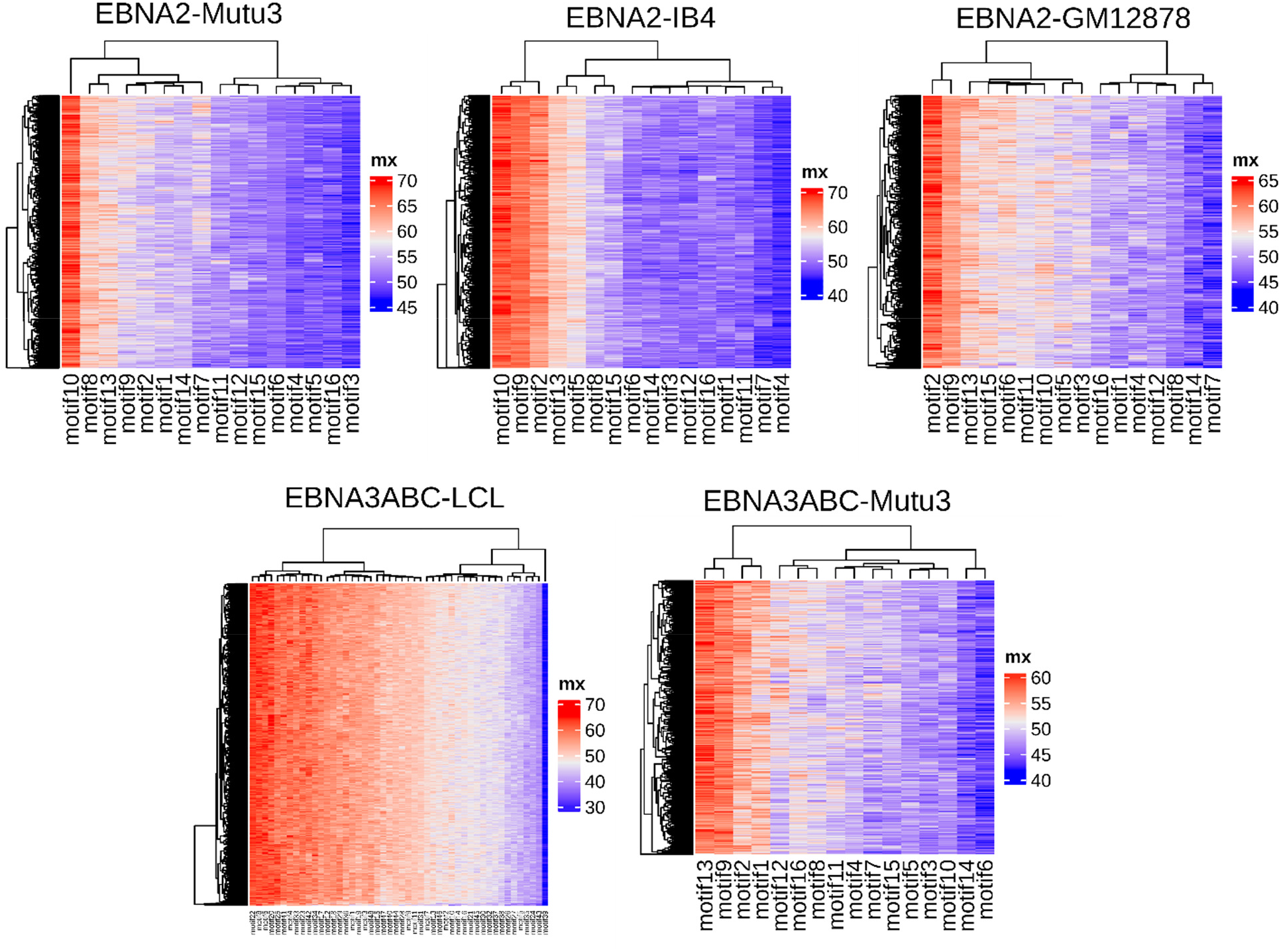
Heatmap showing activation value of each motif on each peak of a dataset. Heatmaps drawn for three EBNA2 cell-lines: IB4, GM12878, Mutu3. And for two EBNA3ABC cell-lines: LCL, Mutu3.

### MuSeAM convolutions reveal both cell-type specific and cell-type independent characteristics of EBNA protein binding

Inter-cell lines motif alignment analysis reveals that some of the co-binding proteins exist in all the three cell-lines, some exist on two cell-lines and some are unique to individual cell-lines (Fig 5). For EBNA2, the motifs common to all GM12878, IB4, and Mutu3 cell-lines are COE1 (EBF1), SUH (RBPJ), ZNF18, IKZF1, and ETS2. Motifs common between Mutu3 and IB4 but absent in GM12878 are IRF2, ELF5, ETV5, ETS1, and ERG. Common between Mutu3 and GM12878, absent in IB4 are MTF1, OTX2, ZNF41, and TBX21. Common between IB4 and GM12878, absent in Mutu3 are ZN770, ZN121, REL, NFKB1, and RFX1. Motifs found only in the Mutu3 cell-line are RUNX1, RUNX3, SPI1, EHF, and STAT2. Found only in IB4 are RFX2, ZN263, RFX3, STAT1, and RELB. And only in GM12878 are RARA, EGR1, MAFB, MXI1, and RARBG (Fig 6 and the S4 and S5 Folders).

**Fig 5.**
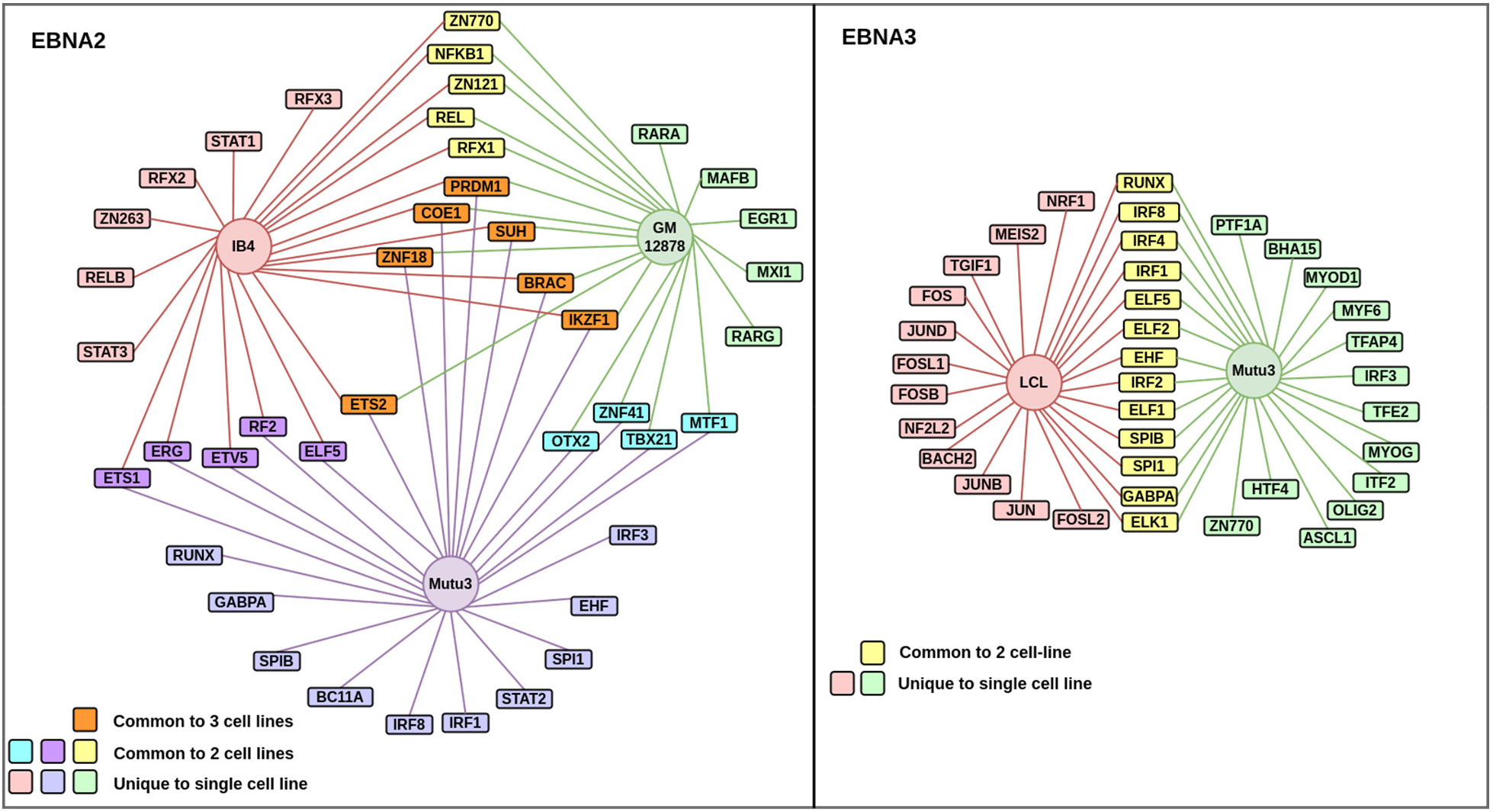
Cobinding network built using the top cell-line specific and cell-line independent TFs of (A) EBNA2 and (B) EBNA3ABC. (For entire network data see the S5 Folder)

**Fig 6.**
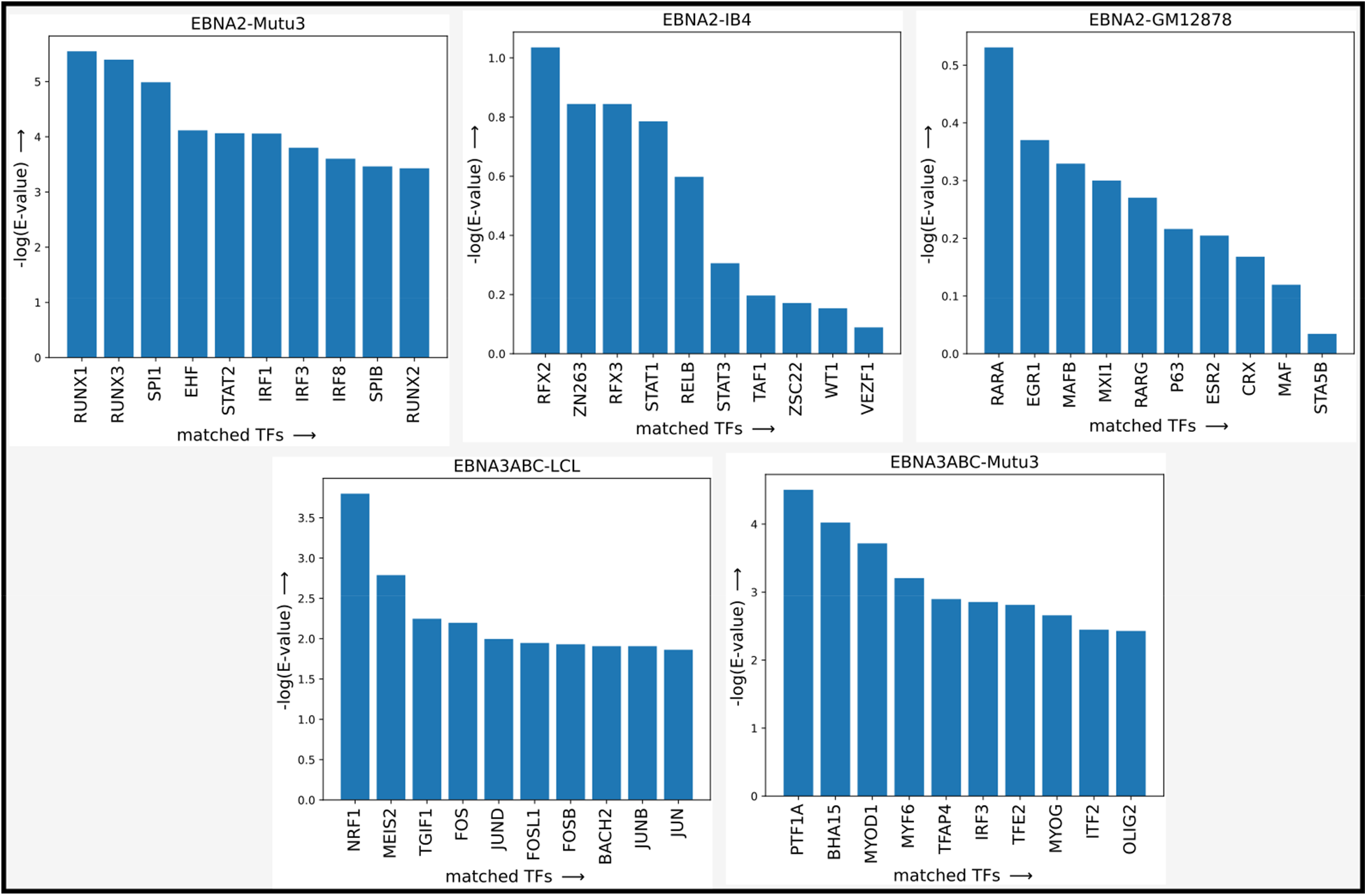
Negative logarithm of E-values of cell-line specific cobinding sites matching with known human motifs.

Motifs common between EBNA3ABC-Mutu3 and EBNA3ABC-LCL are RUNX, IRF8, IRF4, IRF1, and ELF5. Motifs found only in EBNA3ABC-LCL are NRF1, MEIS2, TGIF1, FOS, and JUND. And found only in EBNA3ABC-Mutu3 are PTF1A, BHA15, MYOD1, MYF6, and TFAP4 (Figs 5 and 6).

This lack of common motifs across cell-lines can be partly explained by the fact that EBNA2 ChIP-peaks are largely cell-line specific: The Jaccard index between peaks of the EBNA2 cell-lines are 0.1 (Mutu3 and IB4), 0.097 (Mutu3 and GM12878) and 0.182 (IB4 and GM12878), and between the EBNA3 cell-lines is 0.062 (LCL and Mutu3). However, this also implicates the interesting possibility that vTFs extensively change their co-binding partners in a cell-line specific manner.

### Gene ontology analysis reveals cell-line specific and cell-line independent enriched pathways and processes impacted by EBNA proteins

Similar to the analyses on motifs and co-binding proteins that are cell-line specific and consistent, we next investigated the extent to which the biological processes impacted by EBNA proteins are cell-line specific or consistent across cell-lines. To this end, we first used the tool Homer [30] to identify the genes potentially regulated by the peaks of every vTF. We next analyzed these genesets for enriched gene ontology (GO) terms.

Our analyses revealed that EBNA2 occupied regions potentially impact the immune system and cellular contact related biological processes in all three cell-lines. The processes include leukocyte activation, regulation of cell adhesion, negative regulation of immune system process, and immune effector processes (Fig 7a). We also found processes that are enriched in two cell-lines. For example, protein phosphorylation and cell morphogenesis related processes are more enriched in Mutu3 and IB4, but not in GM12878. Similarly, signal transduction by growth factor receptors and second messengers and regulation of I-kappaB kinase signaling/NF-kappaB signaling related processes are enriched in Mutu3 and GM12878, but not in IB4. Finally, processes related to endocytosis, regulation of proteolysis, and cell death regulation are enriched exclusively in the Mutu3 cell-line.

**Fig 7a:**
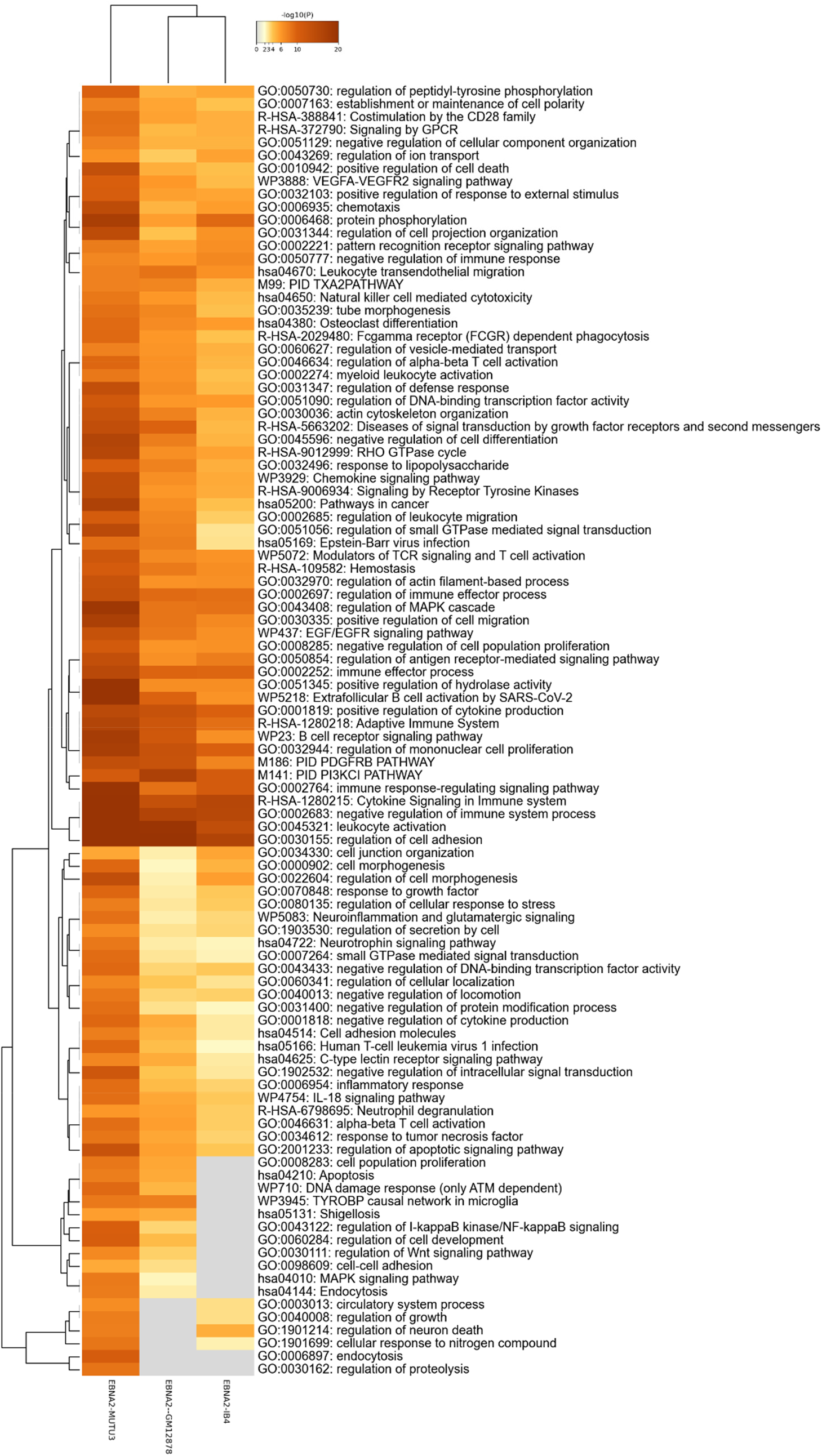
Heatmap of enriched terms across input gene lists colored by p-values for EBNA2.

A similar analysis for EBNA3 proteins on Mutu3 and LCL cell-lines also suggests that some of the pathways and processes impacted are cell-line specific and some are cell-line independent. The GO processes of leukocyte activation, cellular response to cytokine stimulus, regulation of GTPase activity, and regulation of antigen receptor-mediated signaling pathways are common to both LCL and Mutu3 cell-lines. However, the E2F pathway and tumor necrosis factor-mediated signaling pathway are only enriched in the LCL cell-line. On the other hand, tissue morphogenesis, neuroinflammation, and glutamatergic signaling, and regulation of B cell activation are only enriched in the Mutu3 cell-line.

**Fig 7b:**
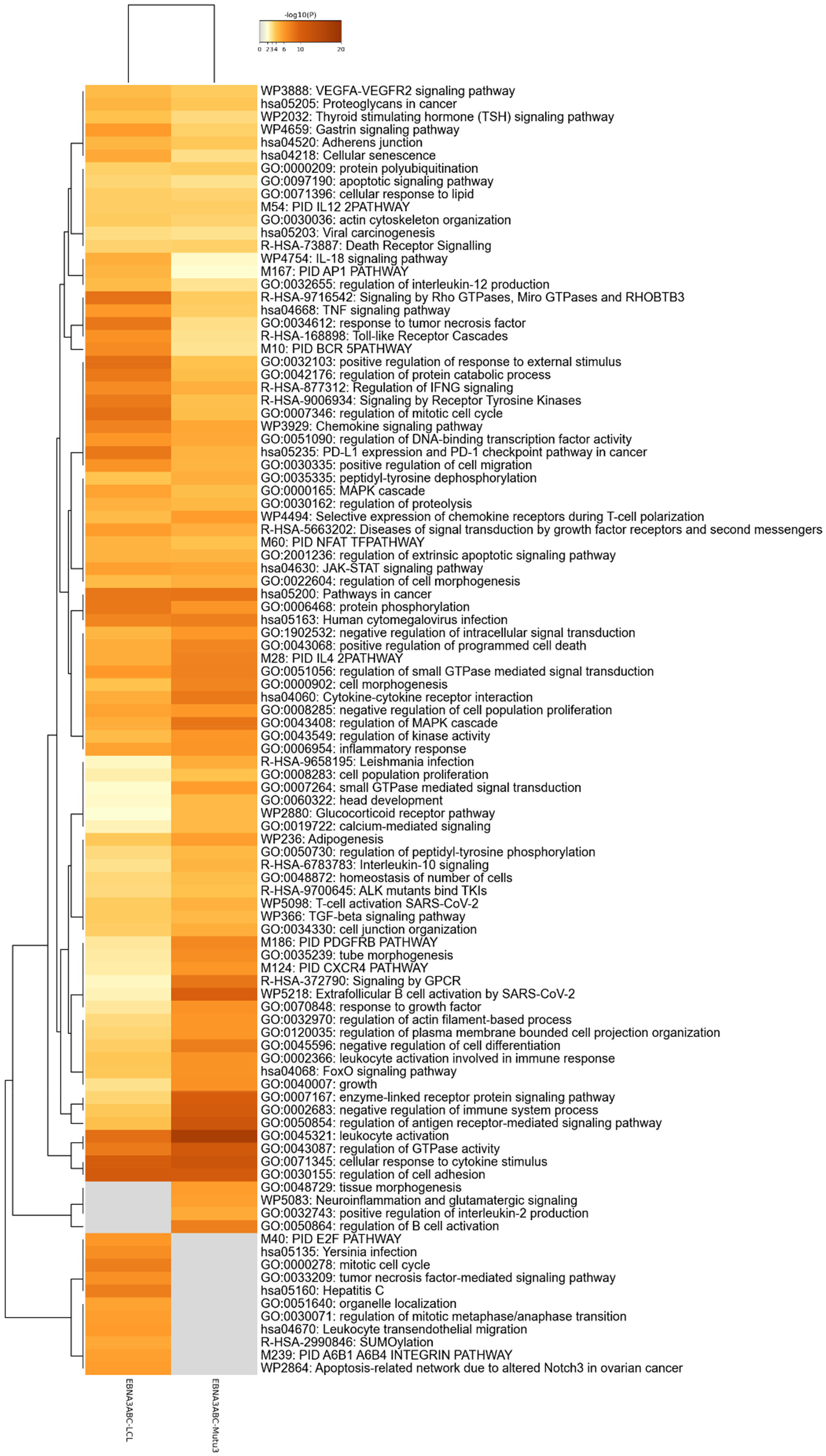
Heatmap of enriched terms across input gene lists colored by p-values for EBNA3.

## Discussion

Using a uniformly processed compilation of ChIP-seq datasets and a state-of-the-art convolutional neural network model for *de novo* motif discovery from genomic sequence data, we performed a systematic analysis of EBNA protein binding specificity in human cell-lines. The key motivation of this study was to discover the “lexicon of sequence motifs” at viral TF binding sites in the human genome, as has been done before for human TFs [31].

Using a *de novo* motif discovery tool enabled us to overcome the potential shortcoming of only testing for the enrichment of known human binding motifs. Indeed, we discovered novel binding motifs at the ChIP peaks of EBNA proteins that do not match with any known motif for human TF binding. This suggests that the EBNA proteins can either directly bind to genomic loci without recruiting cofactors or that some cofactors (from the viral or human proteins) of EBNA proteins are still to be characterized.

Furthermore, we found that both EBNA2 and EBNA3 have the common characteristic of deploying cell-line independent and cell-line specific binding mechanisms. To our knowledge, distinguishing these two mechanisms has not been systematically studied before. However, our study highlights the need to consider both mechanisms when investigating the therapeutic vulnerabilities of these proteins. We also find that both proteins impact some biological processes that are common across cell-lines, but also processes that are exclusively impacted in one or a few cell-lines. This latter observation suggests that these proteins can lead to cell-line specific reorganization of protein-protein interactions, another critical aspect to consider during drug design.

Overall, our study laid out a roadmap for future integrative analysis of vTFs, distinguishing how their DNA binding mechanisms vary across tissues and discovering potential approaches for effective drug design against these viral infections.

## Methods

### ChIP-Seq raw data preprocessing

We used a total of seven datasets (three for EBNA2 and one for each of EBNA3A, EBNA3B, EBNA3C, and EBNA3ABC) mentioned the review of vTFs by Liu et al. [18] GEO links to the datasets are as follows. EBNA2-IB4 - https://www.ncbi.nlm.nih.gov/sra?term=SRP00787

EBNA2-GM12878 - https://www.ncbi.nlm.nih.gov/geo/query/acc.cgi?acc=GSM2039170

EBNA2-Mutu3 - https://www.ncbi.nlm.nih.gov/sra?term=SRX290877

EBNA3ABC-Mutu3 - https://www.ncbi.nlm.nih.gov/geo/query/acc.cgi?acc=GSM1153766

EBNA3A-LCL - https://www.ncbi.nlm.nih.gov/geo/query/acc.cgi?acc=GSE76166

EBNA3B-LCL - https://www.ncbi.nlm.nih.gov/geo/query/acc.cgi?acc=GSE76166

EBNA3C-LCL - https://www.ncbi.nlm.nih.gov/geo/query/acc.cgi?acc=GSM1975602

To preprocess the datasets, we followed ENCODE’s recommendations for ChIP-seq data regarding genome indexing, fastq concatenation, alignment, and filtering. The details are available at: https://www.encodeproject.org/about/experiment-guidelines/

The resulting narrow peak formatted datasets were converted into bed formats using a custom bash script that copies the first six columns of a narrow peak formatted file and outputs as a bed file.

### Generating negative sequences

For generating the negative sequences for each of the datasets, we used the gkmSVM R package. This package imports the gkmSVM v.20 functionalities into R and uses the kernlab library for various SVM algorithms [32]. This package provides an implementation of a new SVM kernel method using gapped k-mers as features for DNA or protein sequences. But we only used the genNullSeqs function of the package. This function generates null sequences (negative set) with matching repeat and GC content as the input bed file for positive set regions. It takes positive set regions which is a bed file, genome version, xfold which is the count of the null sequences, tolerance for difference in repeat ration, tolerance for difference in GC content, batch size, the maximum number of trials, and genome as input arguments. We used the genome version of ‘hg19’. The xfold value was initially set to default value 1 which indicated the same number as in the positive set. But sometimes gkmSVM failed to generate an equal number of sequences and so we set the value greater than the actual length of the positive set so that even if it fails to generate the given number of negative sequences, the overall count of the null set should be greater than or equal to the count of the positive set. The value of tolerance for difference in repeat ratio was the default value of 0.02. The value of tolerance for difference in GC content was also the default value of 0.02. The batch size was also the default value of 5000. The maximum number of trials argument was set to the default value of 20. The genome argument was set to the default value of null. It writes the null sequences to the file with the provided filenames. It actually gives 3 output files. One is for the null sequence genomic regions in bed format, another is for the positive set sequences in the fasta format and the other is for the negative set sequences in the fasta format. We used both the positive and negative fasta formatted output files.

### Preprocessing dataset

The positive and negative sequences of fasta formatted files were merged together into a single file and we generated an output file for the merged file. We stacked the positive sequences over negative sequences in a single file in order. The number of positive sequences and negative sequences was the same. The output line contains 1 for a positive sequence and 0 for a negative sequence in each row for the corresponding sequence in the merged file. So, the total number of rows of the output file is the same as the total number of sequences in the merged file. We used a python script for these purposes.

As the positive and negative sequences were in order, they were randomly shuffled for each dataset. Then for each sequence, the reverse complement sequence was generated. For this, we first complimented the sequence by replacing ‘A’ with ‘T’, ‘C’ with ‘G’, ‘G’ with ‘C’, and ‘T’ with ‘A’.Then this complemented sequence was reversed to generate the corresponding reverse complement sequence.Then both the original sequence and the complemented sequence were transformed into an integer encoded sequence where ‘A’ was replaced by 0, ‘C’ by ‘1’, ‘G’ by 2, and ‘T’ by 3. Then the integer encoded sequences were transformed into one hot encoded sequence where the two dimensional input became three dimensional input by replacing ‘0’ with ‘1000’, ‘1’ with ‘0100’, ‘2’ with ‘0010’, and ‘3’ with ‘0001’.

The length of each sequence in a dataset was different. So to make the length of all the sequences of a dataset equal, we padded ‘0000’ the required number of times to make the sequence length the same as the maximum length of the sequence among all the sequences of the dataset. In a similar way, the labels of each sequence that are in the output file were also converted to a one hot encoded sequence by replacing ‘0’ with ‘10’ and ‘1’ with ‘01’.

Then for 10-fold cross validation, the transformed dataset was split into 10 folds in a stratified way so that the ratio of positive and null sequences are equal in each fold. Every fold has an equal number of sequences.

### Learning CNN model

We used both a custom CNN called MuSeAM and a conventional CNN to learn the models for each dataset. In the conventional CNN, the input layer is followed by a 1D convolution layer. The size of the input layer differs from dataset to dataset. Both the forward one hot encoded sequences and reverse complement one hot encoded sequences were passed to the convolution layer independently having the same weight matrix. Then the two output layers were concatenated. This concatenated output layer was followed by a max pooling layer. Then a flatten layer was applied which flattens the output of the max pooling layer. After that, a fully connected linear layer was applied which was followed by a ReLU activation layer. Finally, a classification layer, which is a fully connected linear layer having 2 features followed by a softmax layer, was applied.

On the contrary, the MuSeAM model used a custom 1D convolution layer instead of the conventional convolution layer as indicated in the MuSeAM model. The rest of the architecture was the same as the conventional CNN architecture.

We then fitted the model using 10-fold cross-validation and tuned the hyperparameters of the model. To meet the overfitting problem, we used L1 regularization. We used accuracy and AUROC as metrics. Finally, we retrained the model using the tuned hyperparameters on the full dataset and computed AUROC and accuracy on the full dataset. The tuned hyperparameters values, model accuracy, and AUROC are provided in the S1 Table.

### Benchmarking learned models

All the models learned using MuSeAM and conventional CNN were compared against each other. We compared the AUROC and accuracy of each model performing on the dataset of other cell lines of the same viral protein. The model learned on a dataset of a cell line of EBNA2 viral protein was tested on the other datasets of cell-lines of EBNA2 viral proteins. Similarly, the model learned on a dataset of a cell line of an EBNA3 viral protein was tested on the other datasets of cell lines of EBNA3 viral proteins. We tested using both the architecture models and compared the AUROC and accuracy.

### Generating motifs

After training the model on a dataset, we generated motifs from the first (and only) convolution layer. The motifs we generated can be thought of as a sequence of probability sets, each set containing the probabilities of every nucleotide (A, C, G, and T for DNA). So if we generated a motif of length 10, then there would be 10 probability sets, each containing a probability for all four nucleotides. So in total, there would be 40 probabilities.

The process worked as follows, from each kernel/filter of the convolution layer we generated a single motif. Each of our kernels can be thought of as a sequence of activation value sets, each set containing an activation value for every nucleotide. So we took each of these activation value sets (a set of 4 activation values), multiplied them by ɑ (a hyperparameter), subtracted from their maximum value, and finally applied softmax to get a probability set. So from each activation value set of our kernel, we got a probability set, giving us a motif. We stored these motifs in a .*meme* file. We then drew these motifs and made logos from them. The background probability used for nucleotides ‘A’ and ‘T’ was 0.3 and for ‘C’ and ‘G’ was 0.2. We used Shannon’s information content with uniform background probability to draw the motifs.

### Identification of human gene binding sites

After generating the motifs, we used the Tomtom [23] tool to compare our generated motifs with the human genome library. The “-no-ssc” option was used for logo generation in the comparison. The motifs were allowed to be reverse complemented before comparison, the minimum overlap was set to 5, and Euclidean distance was used as the distance measure. The maximum E-value threshold was set to both 1 and 10, and a full set of comparison results were generated with both thresholds. Two separate sets of comparison results were generated for the HOCOMOCO-v11 Core Human genome library [33] and the HOCOMOCO-v11 Full Human genome library. The Tomtom tool produced HTML, TSV, and XML results from which we saw the similarities between our generated motifs and the already known human genomes that partake in biological pathways.

### Inter cell-line motif comparison

We used the Tomtom tool to generate comparison results between pairs of our generated motifs of the same vTF in different cell-lines. The comparison options were the same as when comparing with the known human genome library. The comparison pairs were:

- EBNA2-GM12878 vs EBNA2-IB4
- EBNA2-Mutu3 vs EBNA2-IB4
- EBNA2-GM12878 vs EBNA2-Mutu3
- EBNA3ABC-Mutu3 vs EBNA3ABC-LCL

Then we used the tsv results of each dataset when comparing with the known human genome library and used set operations to find the genes common to different combinations of vTFs and cell-lines. For the three cell-lines (IB4, Mutu3, and GM12878) of EBNA2, we found the genes that were common between all of them (meaning these genes were predicted to be present in all cell-lines by our model), genes that were common between only two of the cell-lines, and genes that were present in only one of the cell-lines. We did the same for EBNA3ABC.

### Motif importance ranking

After training the models and generating the motifs we needed to rank the motifs on some score to try to understand their importance and to set a threshold of how important a motif needed to be before we could call it a novel. Our first attempt at this was to take the kernels corresponding to each motif and turn them off or remove them from the convolution layer and then run the model on the dataset again. If the kernel was indeed important then the metrics (Accuracy, AUROC) should drop after removing it. We did this for each kernel and used the normalized accuracy drop for each kernel as its importance score. Next, we took the E-values of the best matches of each motif and normalized the negative logarithm of these E-values to give them another importance score. Both of these importance scores are in the [0, 1] range since they are both normalized values.

### Motif clustering

We also attempted to find if the motifs were only matched on a subset of the peaks in each dataset. For each motif we found its “Activation Value” with each peak in its dataset. The process of finding the activation value is as follows: we calculate the probability matrix from a kernel similar to the convolution layer of MuSeAM. We then slide the probability matrix along the sequence of a peak and calculate the convolution values and take the maximum value of all sliding positions. This maximum value is the activation value of that motif on that peak. We then made two dimensional heatmaps of the logarithms of the activation values with the kernels on the x-axis and the sequences on the y axis. The heatmaps were made with the ComplexHeatmap [34] tool which also offers clustering of the kernels based on the activation values.

### Proposing novel viral binding sites

First, the predicted binding sites were compared with known human motifs using an E-value threshold of at most 10. Two motifs did not match any known human motifs even at an E-value threshold of 10. We accepted these motifs as “Tier-1 Novel” motifs. Then we ran the same comparisons with an E-value threshold of at most 1. Now, some more motifs did not match any known human TFs. Of these motifs we accepted the ones with a q-value of 0.95 or above as “Candidate Tier-2 Novel” motifs (there were 21 such motifs). We then visually inspected the 21 Candidate Tier-2 Novel motifs to find potential matches with known human motifs and eliminated 18 of them. The remaining three motifs were accepted as “Tier-2 Novel” motifs. We refer to the two Tier-1 Novel motifs and the three Tier-2 Novel motifs together as the “Novel” motifs.

### Calculating inter cell-line dataset similarities

To measure the similarities between two genomic datasets, we used Jaccard distance of bedtools [35]. Jaccard statistic is used in set theory to represent the ratio of the intersection of two sets to the union of the two sets. Similarly, it can be used in genomic intervals. It measures the ratio of the number of intersecting base pairs between two sets to the number of base pairs in the union of the two sets. The bedtools implements this statistic in a slightly modified way by subtracting the length of the intersection from the length of the union. As a result, the value of Jaccard ranged from 0.0 to 1.0 where 0.0 represents no overlap in the genomic region of the two datasets and 1.0 indicated complete overlap in the genomic regions of the two datasets. We used this on each pair of datasets in the cell-lines of EBNA2 and EBNA3.

### Peak annotation

We annotated the peaks of each dataset using HOMER tools [30]. We used the peakAnnotate function of the homer. We gave the bed file of each dataset and it generated an annotated file consisting of 19 columns. These are PeakID, Chromosome, start, end, strand, peak score, focus ratio, annotation, detailed annotation, distance to TSS, nearest Promoter ID, Entrez Id, Nearest Unizene, nearest refseq, nearest ensembl, gene name, gene aliases, and gene type. We used the gene name column of each dataset for gene-set enrichment analysis.

### Gene set enrichment analysis

For gene set enrichment analysis, we used the metascape tool [36]. We generated the associated genes of each dataset and did a comparative analysis on EBNA2 cell-lines and EBNA3 cell-lines. We gave multiple gene lists of Mutu3, GM12878, and IB4 cell-lines and metascape did the analysis and generated a heatmap of enriched terms across gene lists colored by p values.

## Supporting information

All Supplemental Files

## Supporting Information

**S1 Table. Hyperparameters and metrics**. The tuned (and finally selected) hyperparameters used during 10 fold cross validation and then during final training. And the corresponding metrics for each run.

**S2 Table. Cross cell line metrics**. For each viral protein, the models trained on one cell line were used to predict viral inhibition on the other cell lines. The table holds metrics (Accuracy and AUROC) found after such cross comparison.

**S3 Folder. Predicted motifs**. The motifs predicted by the trained models for each cell line. Each dataset folder contains the motif logos and the motif meme files.

**S4 Folder. Tomtom comparison results**. Results from Tomtom with E-value threshold at 1.0 and at 10.0.

**S5 Folder. Viral cobinding network data**. Data used to generate the viral cobinding network.

