## Supplementary material for "Epstein-Barr virus nuclear antigen proteins deploy diverse mechanisms to bind the human genome": All Supplemental Files: tomtom.html

Tomtom Results

[close ]

[close ]

[close ]

[close ]

A link to more information about the query motif.

[close ]

[close ]

The motif preview. On supporting browsers this will display as a motif
logo, otherwise the consensus sequence will be displayed.

[close ]

The number motifs in the target database with a significant match to the query motif.

[close ]

Links to the (up to) twenty target motifs with the most significant matches to the query motif.

[close ]

[close ]

The number of motifs read from the motif database minus the number that
had to be discarded due to conflicting IDs.

[close ]

The number of motifs in this database that have a significant match to at least one of the query motifs.

[close ]

The summary gives information about the target motif. Mouse over each
row to show further help buttons for each specific title.

[close ]

The ID of the target motif with the optional alternate ID shown in parentheses.

[close ]

[close ]

[close ]

[close ]

[close ]

[close ]

[close ]

[close ]

The image shows the optimal alignment of the two motifs. The sequence logo
of the target motif is shown aligned above the logo for the query motif.

[close ]

By clicking the link "Create custom LOGO ↧" a form to make custom logos
will be displayed. The download button can then be clicked to generate a motif
matching the selected specifications.

[close ]

Two image formats, png and eps, are available. The pixel based portable
network graphic (png) format is commonly used on the Internet and the
Encapsulated PostScript (eps) format is more suitable for publications
that might require scaling.

[close ]

Toggle error bars indicating the confidence of a motif based on the
number of sites used in its creation.

[close ]

Toggle adding pseudocounts for **S**mall **S**ample
**C**orrection.

[close ]

Toggle a full reverse complement of the alignment.

[close ]

Specify the width of the generated logo.

[close ]

Specify the height of the generated logo.

[close ]

[close ]

[close ]

|  |  |
| --- | --- |
| Image Type | PNG EPS |
| Error bars | yes no |
| SSC | yes no |
| Flip | yes no |
| Width |  |
| Height |  |

### Tomtom

#### Motif Comparison Tool

For further information on how to interpret these results please access
https://meme-suite.org/meme/doc/tomtom-output-format.html.  
To get a copy of the MEME software please access
https://meme-suite.org.

Query Motifs  |  Target Databases  |  Matches  |  Settings  |  Program information  |  Results in TSV Format 
  |  Results in XML Format

#### Query Motifs

Next Top

| Database | ID | Alt. ID | Preview | Matches | List |
| --- | --- | --- | --- | --- | --- |

#### Target Databases

Previous Next Top

| Database | Used | Matched |
| --- | --- | --- |

#### Matches to

Previous Next Top

| Summary | Optimal Alignment |
| --- | --- |
| Name \|; Database \|; *p*-value \|; *E*-value \|; *q*-value \|; Overlap \|; Offset \|; Orientation \| Show logo download options | ⤒ ↥ ⇘ ↧ ⤓ |

#### Settings

Previous Next Top

###### Alphabet

###### Other Settings

|  |  |
| --- | --- |
| Strand Handling | Reverse complements are not possible so motifs are compared as they are provided. Motifs are compared as they are provided. Motifs may be reverse complemented before comparison to find a better match. |
| Distance Measure | Average log-likelihood ratio Euclidean distance Kullback-Leibler divergence Pearson correlation coefficient Sandelin-Wasserman function Bayesian Likelihood 2-Components score (from 1-component Dirichlet prior) Bayesian Likelihood 2-Components score (from 5-component Dirichlet prior) Log likelihood Ratio score (from 1-component Dirichlet prior) Log likelihood Ratio score (from 5-component Dirichlet prior) |
| Match Threshold | Matches must have a *E*-valueq-value of  or smaller. |

Previous Top

###### Tomtom version

 (Release date: )

###### Command line

  
  
Result calculation took  seconds
